# Comparative Genomics of Firmicutes reveals probable adaptations for xylose fermentation in Thermoanaerobacterium saccharolyticum

**DOI:** 10.1101/2023.03.08.531619

**Authors:** Mateus B. Fiamenghi, Juliana S. Prodonoff, Guilherme Borelli, Marcelo F. Carazzolle, Gonçalo A. G. Pereira, Juliana José

## Abstract

Second-generation (2G) ethanol is one potential biofuel that could be used to achieve the goal of reducing greenhouse gas emissions. Many challenges still need to be overcome for the feasibility of this technology, most of them related to consumption of xylose, a pentose sugar not easily metabolized by industrial microorganisms. Thus, exploring genes, pathways and other organisms that can ferment xylose is a strategy implemented to solve industrial bottlenecks. *Thermoanaerobacterium saccharolyticum* (*T. sac*) is an organism from the firmicutes phylum, capable of naturally fermenting compounds of industrial interest, such as xylan and xylose. Understanding evolutionary adaptations may help not only to solidify this bacterium as a potential substitute to the yeast *Saccharomyces cerevisiae* in industry, but also bring novel genes and information that can be used for yeast, enhance its fermenting capabilities, and increase production of current bio-platforms. This study presents a deep evolutionary study of members of the firmicutes clade, focusing on adaptations that may be related to overall fermentation metabolism, especially for xylose fermentation. One highlight is the finding of positive selection on a xylose binding protein of the xylFGH operon, close to the annotated sugar binding site, with this protein already being found to be expressed in xylose fermenting conditions in a previous study. Results from this study can serve as basis for searching for candidate genes to use in industrial strains or to improve *T. sac* as a new microbial cell factory, which may help to solve current problems found in the biofuels industry.

## 1. Introduction

The need to restructure the global energy matrix and mitigate greenhouse gas emissions has led efforts to find clean alternatives to fossil fuels. One approach is the use of microorganisms for fermentation of lignocellulosic biomasses such as sugarcane bagasse and straw to ethanol, which is particularly interesting as it helps to alleviate competition with foods production. This strategy is known as second generation (2G) ethanol production. The main challenges associated with 2G ethanol are the forming of fermentation inhibitors, such as HMF and acetate, after the feedstock pretreatment step for exposure of its sugars, finding optimal enzymes for breaking sugars into usable monomers, and lack of proper consumption of pentose sugars, such as xylose, contained on these feedstocks by organisms used industrially, such as the yeast *Saccharomyces cerevisiae*.

Xylose metabolism mainly follows two pathways: an oxireductive pathway, comprising a conversion of xylose into xylitol by the enzyme xylose reductase, followed by a conversion of xylitol into xylulose by xylitol dehydrogenase, and finally xylulokinase converts xylulose into xylulose-5-P, which enters the pentose phosphate pathway, ending up in the glucose pathway. In many organisms, including industrial yeast, this pathway has some issues regarding cofactor imbalance on the first two steps, which causes accumulation of xylitol [40]. The other pathway is similar to the first but comprises a single step between xylose and xylulose, done by xylose isomerase, and is predominantly found in bacteria [83]. One bottleneck common to both pathways is related to pentose transport, in which sugar transporters preferentially uptake glucose in detriment of xylose, turning the 2G process unfeasible due to fermentation time increase.

*Thermoanaerobacterium saccharolyticum* (hereafter called *T. sac*) is a bacterium from the firmicutes phylum, found originally in hot springs around Yellowstone National Park [41], capable of naturally fermenting compounds of industrial interest, such as xylan and xylose, and engineered to produce ethanol at higher yields [63]. This organism, among other thermophilic bacteria, has gained some attention over the last decade as an alternative for industrial fermentation, as it grows in higher temperature, similar to those used for pre-treatment enzymes, and can co-ferment both cellulose and hemicellulose sugars [15], a trait seen in many *Thermoanaerobacter* [43,72], which means potential for simultaneous saccharification and fermentation. Also, its ease of transformation confers an additional advantage for metabolic engineering [46,47]. Studies have explored the underlying mechanisms of pentose metabolism in *T. sac*, such as inactivating redox sensing molecules to alleviate alcohol dehydrogenase repression [84], deleting genes that create undesirable byproducts such as acetate and lactic acid [63], discovering essential genes [14], and elucidating fermentation products [32]. However, an evolutionary approach to understand key adaptations to fermentation stresses and describing candidate genes has not been reported for this organism. Evolutionary analyses for better explaining adaptations to industry and suggesting genes for genetic engineering have been successfully used in yeast [7,11,21,57,77]. Understanding its evolutionary adaptations may help not only to solidify this bacterium as a potential substitute to the yeast *Saccharomyces cerevisiae* in industry, but also bring novel genes and information that can be used for yeast, enhance its fermenting capabilities, and increase production of our current bio-platforms.

Regarding again xylose transport, most known transporters are inhibited by glucose or other hexoses, and preferentially carry these sugars instead of xylose during co-fermentation conditions. However, while yeast only have MFS transporters, which are proteins capable of carrying solutes passively through the membrane, bacteria have an additional class called ABC transporters, which are usually organized in operons, that require the use of ATP molecules to pump solutes inward [13,42]. In *E. coli*, the ABC transport system responsible for specifically uptaking xylose is the XylFGH operon, comprised of proteins XylF (external protein responsible for sugar ligation), XylG (ATP-binding) and XylH (translocation of xylose through the membrane), with XylR acting as a regulator. Even though an energy expenditure is needed for consuming xylose, this could confer an advantage when sugar availability is scarce, allowing organisms with these systems to continue thriving in their environment [13,67].

*T. sac* being a lignocellulosic fermenter with low glucose inhibition could have adaptations in its transporters to efficiently carry xylose, specifically, on the ABC transporters because of their energy consumption load. In this work comparative genomics was used across 20 bacterial species, to identify evolutionary marks related to xylose fermentation. Positive selection was found on the xylF homolog, the sugar binding molecule on ABC transporter, where the largest affinity is needed for xylose transport is shown. Together with adaptations in transport proteins, other *T*.*sac* proteins were found to carry important adaptations putatively related to the xylose metabolism. Understanding all these *T*.*sac* adaptations may further help alleviate uptake difficulties on current cell factories such as yeast.

## 2. Materials and Methods

### 2.1 Dataset

20 bacterial genomes, including *T*.*sac*, were downloaded from NCBI for analysis. Genomes were chosen based on proximity to *T*.*sac* (close and distant species), xylose metabolism capacity and industrial applications (table 1).

**Table 1.** Information on the used public bacterial genomes obtained on NCBI

| Bacteria Name | Abbreviation | Xylose metabolism | Assembly ID | Reference |
| --- | --- | --- | --- | --- |
| *Salmonella enterica* | CAD | No | GCA_000195995.1 | [53] |
| *Escherichia coli* | AAC | Yes | GCA_000005845.2 | [6] |
| *Clostridium acetobutylicum* | AAK | No | GCA_000008765.1 | [52] |
| *Bacillus subtilis* | CAB | No | GCA_000009045.1 | [39] |
| *Lactobacillus fermentum* | BAG | No | GCA_000010145.1 | [49] |
| *Lactobacillus acidophilus* | AAV | No | GCA_000011985.1 | [4] |
| *Clostridium botulinum* | ABS | No | GCA_000017045.1 | [64] |
| *Natranaerobius thermophilus* | ACB | No | GCA_000020005.1 | [82] |
| *Ruminiclostridium (Clostridium) cellulolyticum* | ACL | Yes | GCA_000022065.1 | [27] |
| *Thermoanaerobacterium thermosaccharolyticum* | AFK | Yes | GCA_000145615.1 | [27] |
| *Halanaerobium praevalens* | ADO | No | GCA_000165465.1 | [31] |
| *Halanaerobium hydrogeniformans* | ADQ | Yes | GCA_000166415.1 | [9] |
| *Bacillus cellulosilyticus* | ADU | No | GCA_000177235.2 | [48] |
| *Thermoanaerobacterium xylanolyticum* | AEF | Yes | GCA_000189775.3 | [41] |
| *Thermoanaerobacterium saccharolyticum* | AFK | Yes | GCA_000307585.2 | [15] |
| *Thermoanaerobacterium aotearoense* | ETO | Yes | GCA_000512105.1 | [2] |
| *Bacillus beveridgei* | AOM | No | GCA_001721685.1 | [5] |
| *Natranaerobius trueperi* | OWZ | No | GCA_002216005.1 | [24] |
| *Clostridium chauvoei* | SLK | No | GCA_900168365.1 | [60] |
| *Thermus thermophilus* | YP | No | GCF_000091545.1 | Masui et al |

### 2.2 Orthology assignment

Comparing genes across species requires clustering of their proteins into putative homologous groups. Orthofinder [18] was used to accomplish such task, by first clustering genes according to similarity using DIAMOND [10], and cutoff with the default MCL inflation. Briefly, FASTA files of the translated CDS from each species were used as input for DIAMOND, which is used for finding the most similar sequences via reciprocal-best-hits. Orthofinder, by using the MCL parameter, aggregates the most similar sequences into Orthogroups (gene families, a set of genes from multiple species descended from a single gene).Orthogroups of sugar transporter genes were recovered using BLASTP with known protein sequences from *E. coli* or *T*.*sac* and an e-value threshold of 1e-20, and also. through HMM profiles from Pfam (PF07690.15, PF13347.5 and PF05631.13). Genes and protein names were kept as the NCBI identifier. Family names were kept as outputted from Orthofinder.

### 2.3 Phylogenetic inferences

For each gene family, multiple sequence alignment (MSA) was done using protein sequences through MAFFT [34], anchoring first via local alignment (localpair) and 1000 iterations (maxiterate 1000). Phylogenetic relationships for each transporter family’s genes was inferred through Maximum Likelihood, implemented on IQTree [71]. IQTree’s automatic model selection tool was used for fitting the best substitution model and branch support was tested by 1000 ultrafast bootstraps. Species’ phylogenomics was inferred by concatenating all 157 Single-Copy Orthogroups MSAs with FASConcat [37] and analyzing with IQTree using the same strategy as above and through Bayesian methods implemented via MrBayes [58]. MrBayes was ran using a GTR+ invgamma substitution model with 3 hot and 1 cold chains for 10 million generations discarding 25% of the generations from the cold chain

### 2.4 Gene duplication analysis

Gene birth and death estimation was obtained by using the same Orthofinder gene count output and the ultrametric species tree as inputs for CAFE [16] analysis, which models the evolution of Orthogroups based on the species’ phylogeny. CAFE was ran with default parameters, and λ to maximize the log likelihood for all families (lambda -s). Number of expanding (gene birth), contracting (gene loss), average expansion and log of average expansion can be seen on table 2.

**Table 2.** CAFE results for expanding and retracting gene families. EF means number of Expanding Families and CF represents Contracting Families. For both columns, stat means number of statistically significant families (p <0.05)

| Species | EF (stat) | CF (stat) | Avg. expansion | Log Avg. expansion |
| --- | --- | --- | --- | --- |
| *Escherichia coli* | 198 (3) | 226 (1) | 0.00245 | -2.61079 |
| *Salmonella enterica* | 225 (1) | 253 (1) | 0.001685 | -2.77352 |
| *Bacillus beveridgei* | 188 (5) | 845 (2) | -0.09081 | -1.04186 |
| *Bacillus cellulosilyticus* | 293 (5) | 390 (1) | 0.0049 | -2.30976 |
| *Bacillus pseudofirmus* | 338 (2) | 780 (0) | -0.04043 | -1.39331 |
| *Bacillus subtilis* | 380 (4) | 1157 (1) | -0.0781 | -1.10734 |
| *Lactobacillus acidophilus* | 124 (1) | 367 (4) | -0.03277 | -1.4845 |
| *Lactobacillus fermentum* | 108 (5) | 278 (2) | -0.01807 | -1.74303 |
| *Clostridium chaouvoei* | 194 (3) | 1158 (1) | -0.1415 | -0.84924 |
| *Clostridium acetobutylicum* | 308 (7) | 554 (0) | -0.00168 | -2.77352 |
| *Clostridium botulinum* | 279 (3) | 639 (1) | -0.03813 | -1.41871 |
| *Clostridium cellulolyticum* | 283 (11) | 1329 (0) | -0.12021 | -0.92004 |
| *Thermoanaerobacterium xylanolyticum* | 61 (1) | 587 (0) | -0.07672 | -1.11508 |
| *Thermoanaerobacterium saccharolyticum* | 30 (1) | 16 (0) | 0.003216 | -2.49269 |
| *Thermoanaerobacterium aotearoense* | 20 (0) | 25 (0) | -0.00061 | -3.21285 |
| *Thermoanaerobacterium thermosaccharolyticum* | 143 (0) | 579 (0) | -0.06049 | -1.21832 |
| *Natranaerobius trueperi* | 104 (4) | 357 (2) | -0.03247 | -1.48858 |
| *Natranaerobius thermophilus* | 205 (4) | 190 (1) | 0.01853 | -1.73213 |
| *Halanaerobium hydrogenifermentans* | 179 (7) | 345 (1) | -0.01363 | -1.86552 |
| *Halanaerobium praevalens* | 94 (3) | 386 (1) | -0.04196 | -1.37716 |
| *Thermus thermophilus* | 149 (1) | 5330 (0) | -0.79387 | -0.10025 |

Sometimes, a simpler approach can give us interesting insights into genome evolutionary dynamics that may be lost to canonical approaches such as CAFE. Thus, besides this model-based analysis, we carried out a naïve gene duplication analysis by analyzing Orthofinder’s gene count output using a custom python script, by filtering from the dataset gene families where *T*.*sac* had a larger gene count than the mean plus two standard deviations. As retained gene duplications are expected to be scarce, gene families with at least three genes were considered for further detailed examination. Figure 3 shows the results for this analysis.

### 2.5 Natural selection analysis

Nucleotide codon alignments were obtained using MACSE to align the nucleotide sequences by using the protein alignment for each Orthogroup, done through MAFFT (localpair, maxiterate 1000) as template. Codon-based alignments and ML trees for each gene family obtained in OrthoFinder were used in tests for dN/dS models. Selection analysis was implemented using CODEML from the PAML package [80] through ETE3 Toolkit, and MEME from the HYPHY package [51]. Both analyses implement branch-site tests, comparing substitution rates between indicated branches (Foreground) against the other branches (Background). *T. sac* branch tips were marked for both analysis as Foreground for comparison against the other branches. Models ran for CODEML were bsA and bsA1. Positive selection was inferred for sites with p-value of LRT between alternative model and null model (bsA and bsA1, respectively), if the p-value was smaller than 0.05 the site was retrieved as a positively selected site [81]. Families with positive selection detected by CODEML were subjected to MEME to check if a second method would corroborate the CODEML results. The selected sites were retrieved directly by looking if there were branches under selection at a p-value smaller than 0.05.

### 2.6 Functional annotation

Families with positive selection evidence from MEME and CODEML or with duplications were retrieved for further inquiries and discussion. Functional annotation was retrieved from each protein’s NCBI accession, and each family was annotated manually based on the most represented annotation within all proteins. All families with selection were also submitted to Eggnog [30] and PANNZER2 [69] servers as an attempt to better annotate cases in which the original annotation from NCBI was too vague. Briefly, Eggnog uses precomputed Orthogroups with functional description from its database to retrieve the most likely functional information from the inputted sequences. PANNZER2 similarly searches for homolog sequences in the Uniprot database and based on the most similar sequences annotates the most probable function. The most represented annotation within proteins was kept as the family’s annotation.

### 2.7 Microarray analysis

Currie *et al* [15] published microarray data from *T. sac* fermentation on a variety of conditions, including co-consumption of glucose and xylose, and shocking with hemicellulose (which they called ‘washate’). This available data was filtered for proteins of interest obtained from our evolutionary analyses and plotted as a heatmap to help visualize genes with higher expression on these conditions.

## 3. Results

### 3.1 Orthology assignment and phylogenetic inferences

Orthofinder clustering resulted in 14,714 gene families with 2292 families containing at least one T. sac member. Orphan genes (genes not assigned to any family) varied from 1 to 20% for species used (Supplementary figure 1) showing that clustering was satisfactory. A higher orphan rate for some species than others can be explained by the wide phylogenetic range of genomes chosen for analysis.

*T. sac* gene copies for each family were aligned to check if each copy may have diverged since duplication. All sequences showed nucleotide substitutions in alignment, some being much more fragmented, which might indicate pseudogenization, while others maintained much of their structure showing only single nucleotide substitutions.

Both Bayesian and Maximum Likelihood inferences using 159 single copy gene families reconstructed the same relationships among species (figure 1). One interesting finding is that *T. sac* and *T. aotearoense* had no branching between them and showed no nucleotide differences for the 159 single copy genes, which indicates that they are most likely the same phylogenetic lineage and the same species. A reasonable explanation for why this result differs from what is described in literature is because taxonomic classification of these bacteria was done through 16S [45], which is known not to be always a reliable marker for separating closely related species. A newer revision on GDTB [54] also considers *T. aotearoense* as *T. saccharolyticum*.

**Fig. 1.**
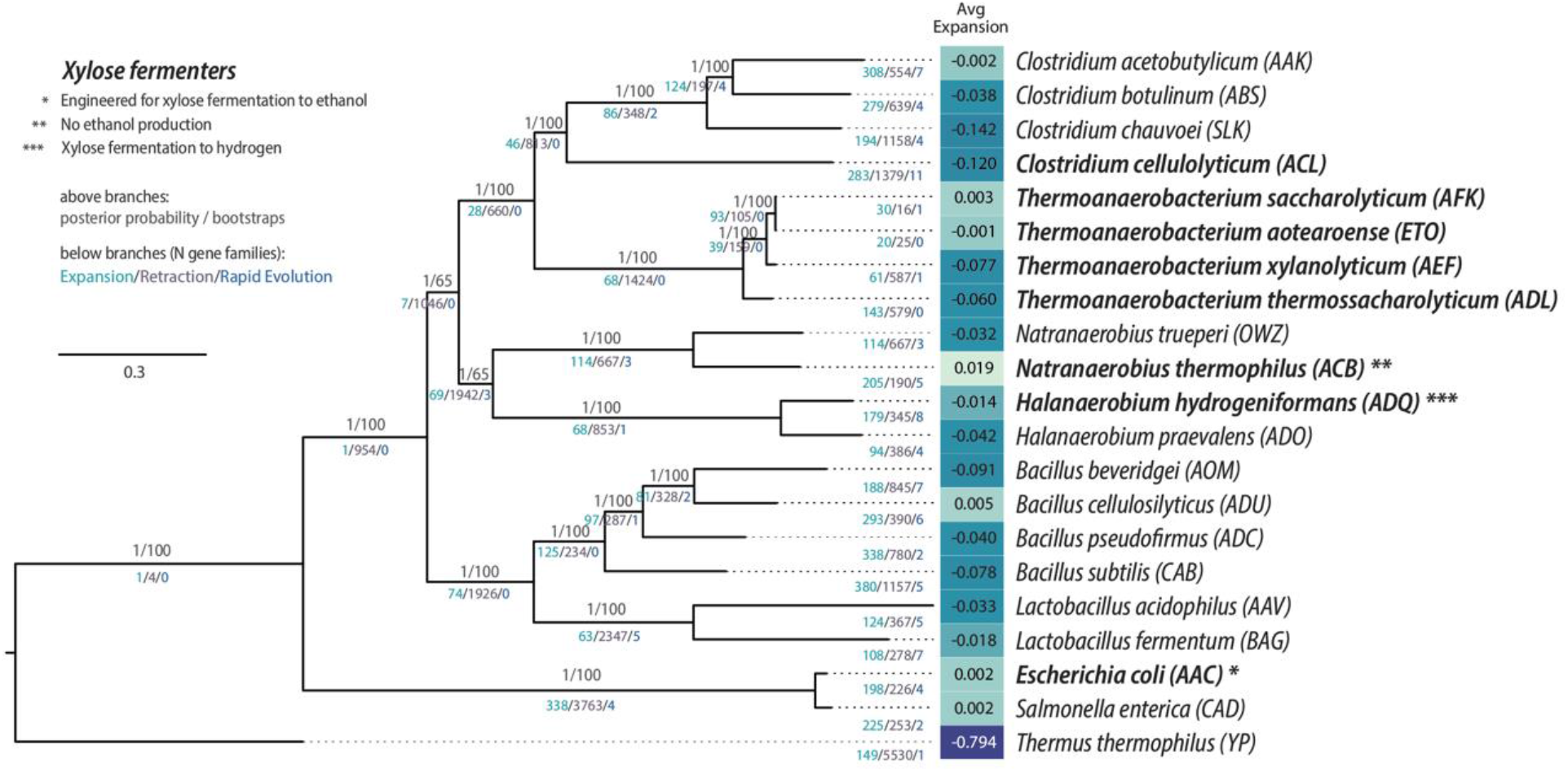
Phylogenetic inference of the 157 single-copy orthogroups for the 20 Firmicutes with *T. thermophilus* as outgroup, through Maximum Likelihood and Bayesian approaches coupled with gene birth and death results. Numbers above branches represent posterior probabilities and bootstraps from MrBayes and IQTree, respectively, and below branches the number of gene families under expansion, retraction or rapid evolution as reported by CAFE. Average expansion was also outputted by CAFE analysis, darker colors represent less expansion (more retraction) while lighter colors represent positive or near zero expansion. Abbreviations after species names represent their NCBI abbreviation which are also used on other figures. Species in bold are able to ferment xylose.

### 3.2 Genome-wide evolution and adaptation clues

The approach of estimating gene birth and losses throughout the genome showed that, on average, we have a tendency for gene losses for all genomes with no relationship to the xylose fermentation phenotype (figure 1 and table 2). Despite almost no expansion seen for T. sac, table 2 shows one statistically significant expansion, which is a rapidly evolving family, found to be OG55, annotated as a transposase IS116/IS110/IS902 family protein. When analyzing *T. sac* family members’ positions in the genome, AFK85526.1 and AFK85527.1 were of special interest, as they appear close to each other, with AFK85525.1 (annotated as a Xylose isomerase domain protein) and AFK85528.1 (annotated as araC) as the closest genes, which are related to pentose metabolism. Besides, this transposase also showed sites under positive selection, as seen both in MEME and CODEML (figure 2).

**Fig. 2.**
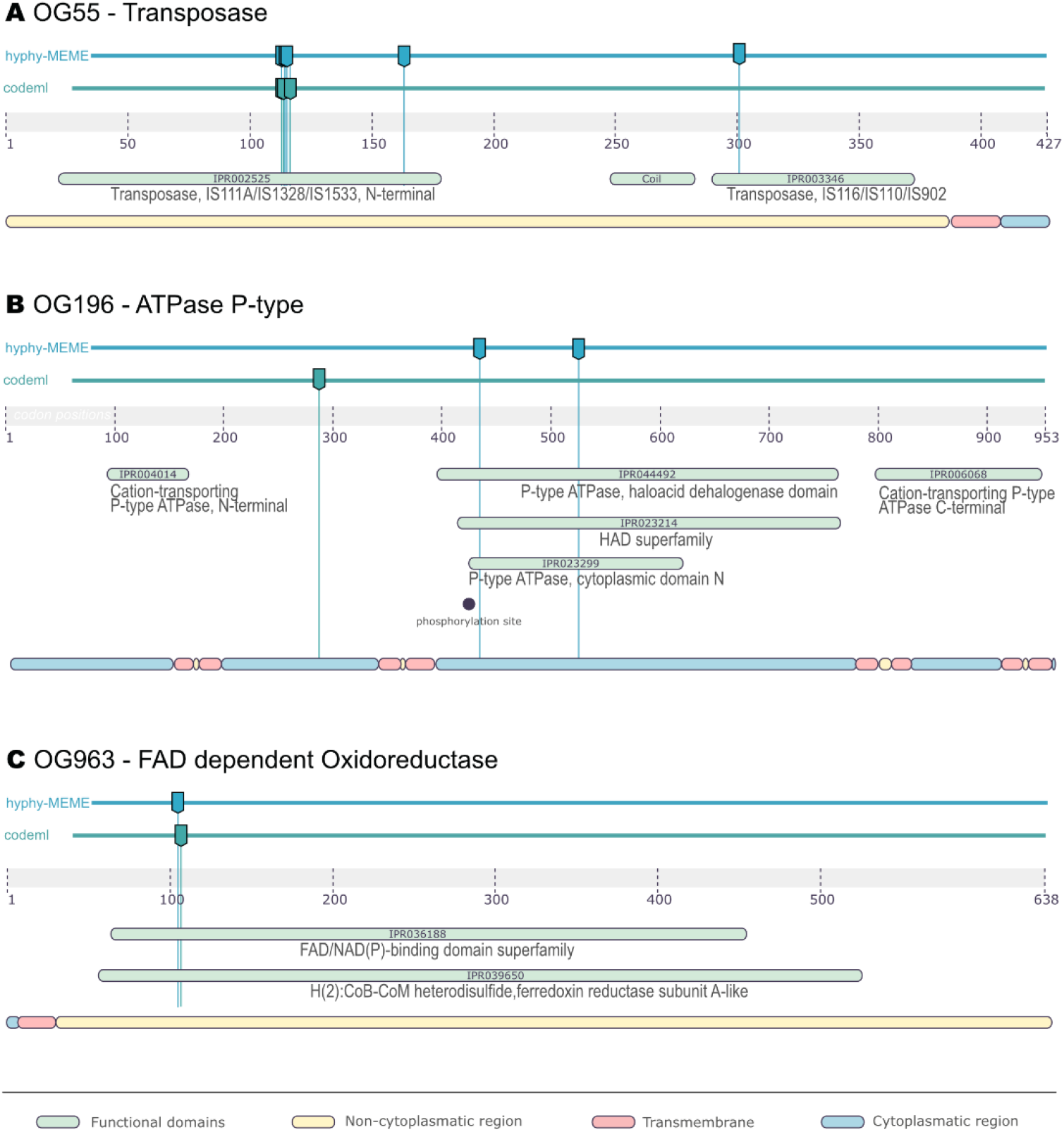
Graphical representation of a *T. sac* protein sequence from families (a) OG55, (b) OG106 and (c) OG963, respectively, with Interpro predicted sites and domains, and sites under positive selection as predicted by HYPHY-MEME and CODEML

To better understand this family’s dynamics with these C5 sugar related proteins, we concatenated all xylose isomerase domain families and reconstructed their phylogeny, which revealed monophiletic clades for the proteins of families OG1593, OG2274, and OG2848, with some mixed clades for OG1258, OG1843 and OG4121 (supplementary figure 2). Xylose isomerase families showed no gene expansion in *T*.*sac*, high sequence conservation, and no positive selection evidence. Both AFK85525.1 and AFK85528.1 were found to have a baseline expression on public microarray data of *T. sac* on xylose (figure 5). Additionally, AFK85525.1 was found in OG4121, a small family containing only *T. sac, T. aotearoense* and *Ruminiclostridium (Clostridium) celullolyticum*, three xylose consuming species.

Regarding positive selection analysis, there were 7 gene families under selection found by CODEML, but only 4 were detected by MEME and thus, we chose to further explore only these four families with coinciding signals: OG55, OG106, OG963 and OG1742 (detailed at a later section). Figure 2 shows OG55, OG106 and OG963 sites with evidence of positive selection, as well as InterproScan [8] predicted domains. The three families showed sites with positive selection within protein domains. Family OG963 was annotated as fumarate reductase/succinate dehydrogenase. Family OG106 comprises HAD superfamily P-type ATPases (PMCA), with a role in translocating calcium, sodium and hydrogen ions across the plasma membrane (Type II subfamily [12]), reducing environmental stress caused by these ions; some downregulation was seen on the microarray xylose conditions for the genes in this family.

### 3.3 Genes possibly related to sugar metabolism with increased gene copies

*T. sac* xylose and glucose co-fermenting phenotype is remarkable and much desired for industrial applications, thus having a bigger viability and fitness against competition, while resisting abiotic stresses, is essential. One interesting family that was found in the analyses was Peptidoglycan Binding Domain 1 (OG1496), which had more copies when compared with other species in this family through the naïve gene duplication approach (3 copies for *T. sac* against the mean of 0.53 copies for the family). This family is involved with cell wall degradation, such as autolytic lysozymes or cleaving autopeptidases, which may suggest a fluidity and dynamic modulation of *T*.*sac* cell wall to resist different stresses [17,22,61].

Another interesting family that followed a pattern of greater copies in *T. sac* than other species was PH1107 glycosidase (OG1224), related to degradation of glycoside bonds of carbohydrates, such as xylan and other complex sugars, which can help to explain its adaptations to ferment these higher carbohydrates directly, even more so with family member AFK87323.1 being slightly upregulated on all conditions when compared to other members of this Orthogroup.

Acetoin dehydrogenase regulator family (OG1421) was surprising as having 5 copies in *T*.*sac*, while the family mean number of copies was 0.62. These genes are responsible for the regulation of acetyl-CoA and acetaldehyde through the reaction acetoin + CoA + NAD^+^ ⇌acetaldehyde + acetyl-CoA + NADH + H^+^ and require thiamine diphosphate as a cofactor. The annotation of a LuxR motif is reported in acetoin regulator for other species, such as *Klebsiella pneumoniae* [28,55].

Pyruvate/Ketoisovalerate oxireductase family (OG1052) was interesting, especially *T. sac* protein AFK87181.1 and AFK86082.1 as this group was also detected on our naïve duplication approach and these members were shown to be expressed in xylose and xylan fermenting conditions when looking at the microarray data (glucose-xylose and xylose_shock). Pyruvate oxireductase is also called pyruvate:ferredoxin oxidoreductase (PFOR), and is involved in the synthesis of acetyl-CoA from pyruvate and oxidized ferredoxin [23,26].

Regarding other sugars, family OG1051, annotated as an extracellular solute-binding protein for the ABC operon MsmEFGK, was interesting. In *Streptococcus mutans* MsmEFGK is responsible for carrying Melibiose, Raffinose, Stachyose, Isomaltose and Isomaltriose [74]. This high copy-number may be explained by a need to transport different solutes, and as an ABC transport requires energy, adjusting these copies to be specific to each sugar may have been what happened during *T. sac* evolution. Also, although we usually see regulation confined on the operon’s region, it has been proposed that different subunits of an ABC transporter can pair with other ABC components to fulfill their function [66,74]. Figure 3 shows information retrieved from our naïve approach.

Proteins from all these families are shown in the microarray (figure 5), with family OG1224 and OG1421 having some of the most expressed genes for most conditions and family OG1051 being mostly repressed or neutral, which is expected as the members are not directly related to xylose.

### 3.4 Genes of xylose transporter families

Six families were identified as MFS transporters and 7 as ABC transporters. The known xylose MFS transporter from *E. coli* xylE was positioned on OG271 according to our BLAST search, however, no *T. sac* genes were present in this family. Analyzing the BLAST results, no *T. sac* or other *Thermoanaerobacterium* genes were similar to xylE within our e-value cutoff threshold. The most similar gene was AFK86490.1, with 29.49% identity, annotated as a drug resistance transporter (OG25, which was added for analysis, data not shown). This initial finding suggests that *Thermoanaerobacterium* strategies for xylose transport differ from other groups of bacteria, which may be indicative of their efficiency for 2G fermentation.

OG1742 (xylF family) was recovered upon closer inspection of proteins annotated as sugar transporters in the *T. sac* microarray published by Currie et al [15]. AFK86454.1 had a higher expression during their experiments than other similar extracellular binding proteins or MFS transporters. As this family also was not returned through BLAST with known xylF proteins, this may also suggest a different mechanism or adaptation for active transport of xylose.

No evidence of positive selection was found in MFS transporters. Also, for the ABC xylose transporter operon no xylG, xylH or xylR families had any evidence of positive selection. However, as also previously reported for OG55, OG106 and OG963, OG1742 (xylF) showed evidence for positive selection for both CODEML and MEME analyses. Figure 4 shows the family’s phylogeny, part of the MSA and sites under positive selection. One interesting finding is that residue 274 (239 ungapped) is two residues distant of site 272 (237 ungapped), which is annotated as one of the sugar binding sites when analyzed through InterproScan.

**Fig 3.**
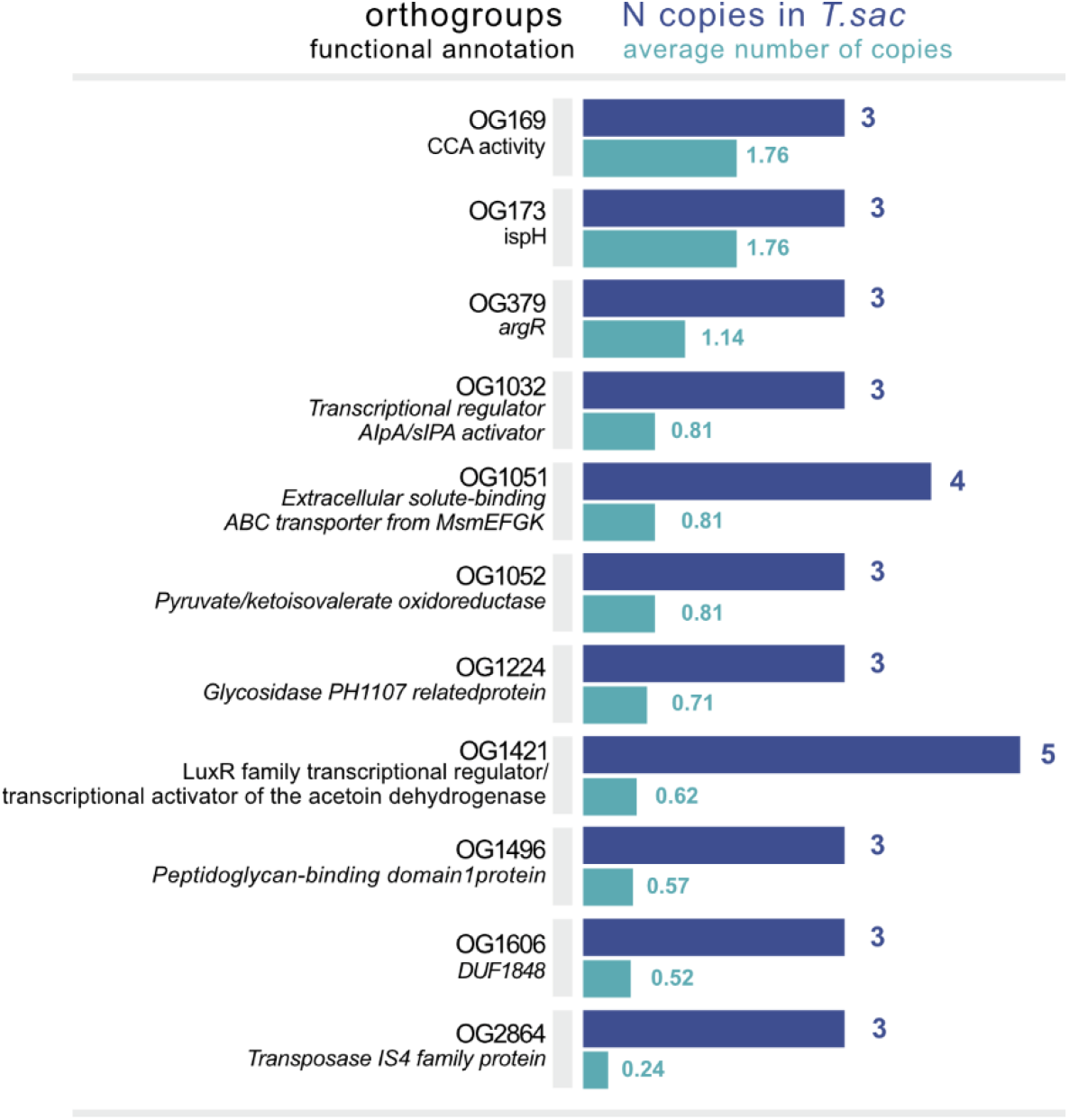
Naïve duplication analysis for gene families in which T. sac has more copies than the family mean plus two standard deviations number of copies, and with at least 3 duplications in *T*.*sac*. Shown in purple are the copies in *T*.*sac* and in green the mean number of copies for the family.

**Fig 4.**
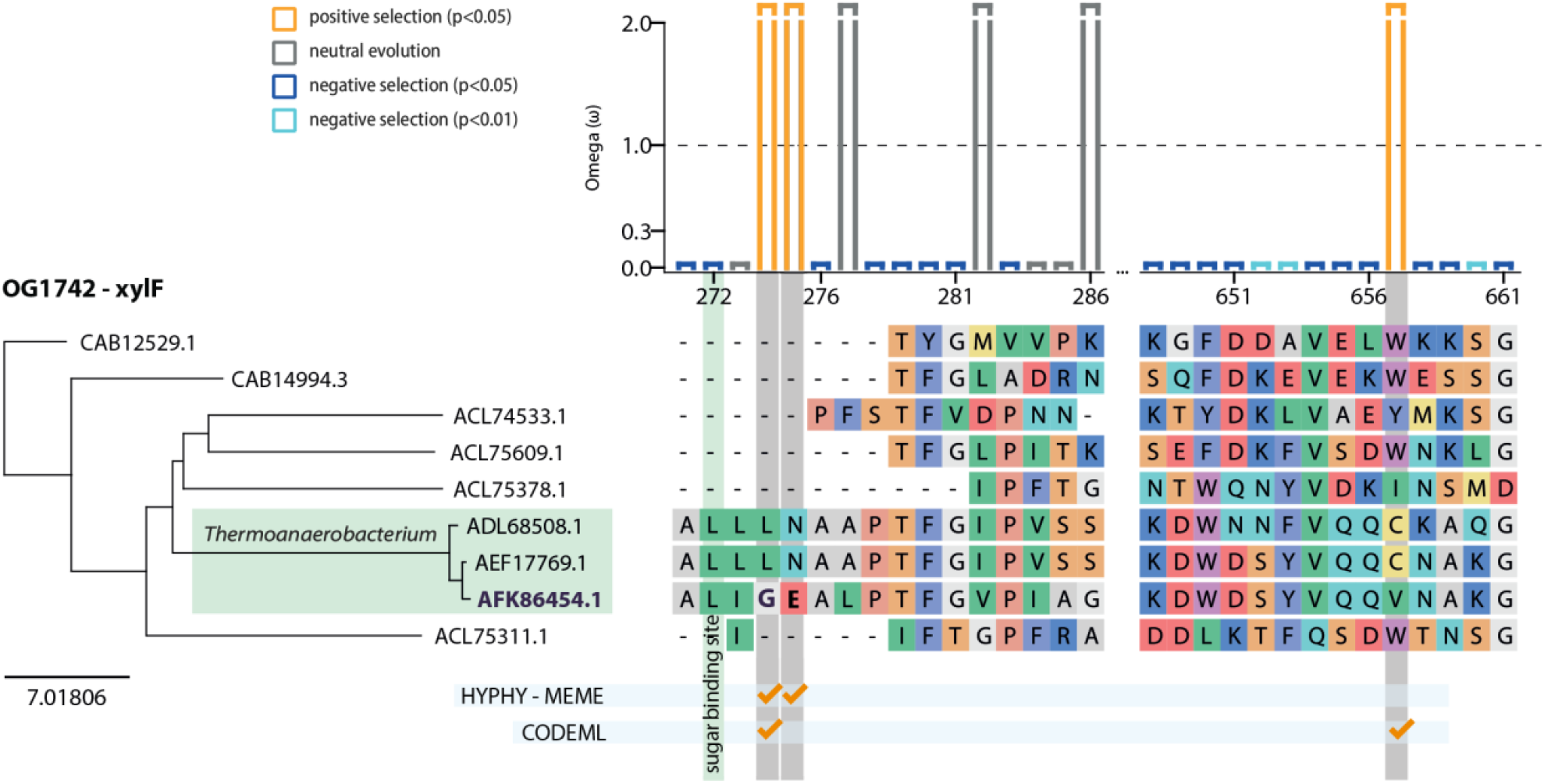
Phylogenetic inference, MSA and sites under positive selection for xylF family OG1742.

**Fig 5.**
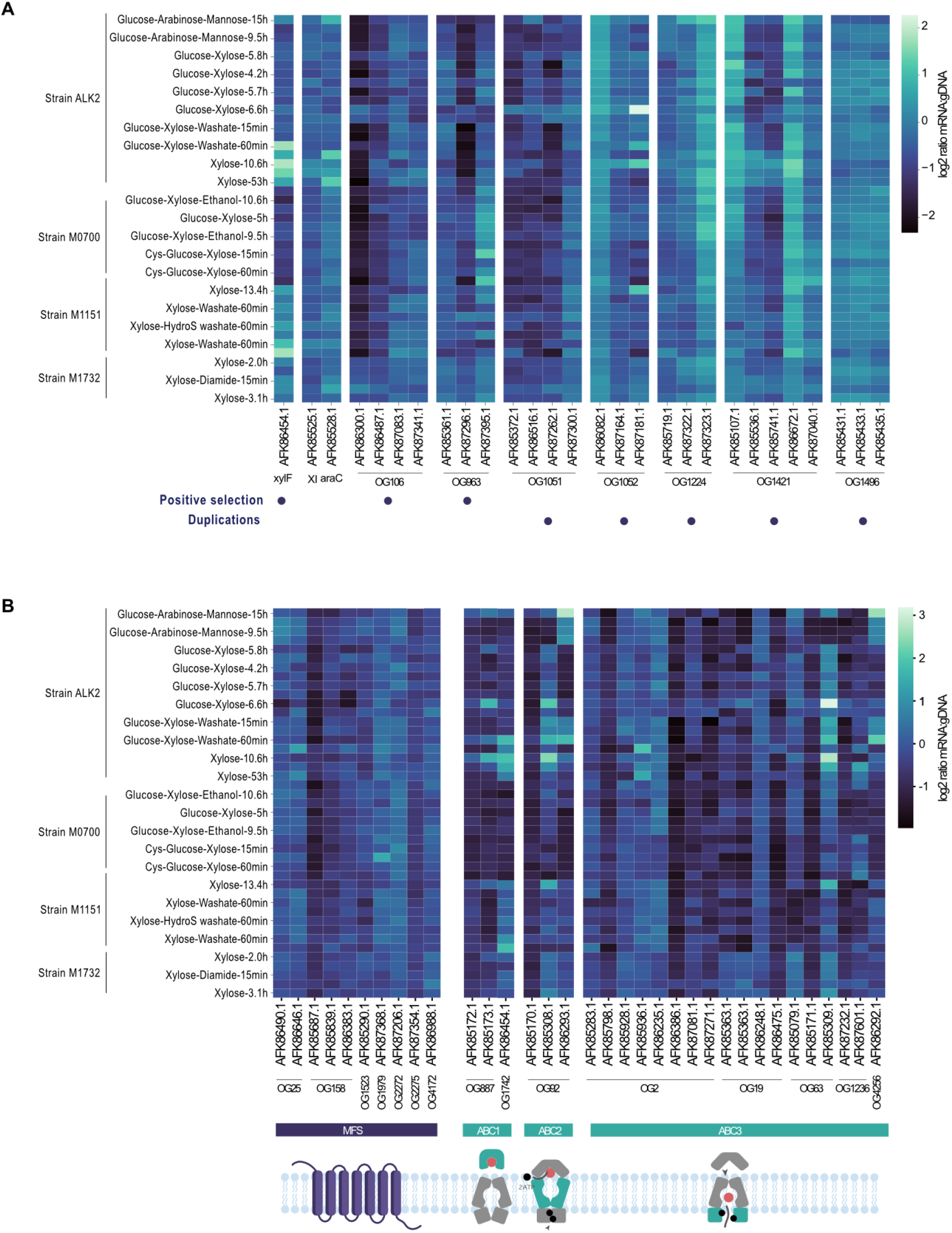
Heatmap adapted from Currie et al microarray data showing (a) xylose related genes with putative adaptations and (b) sugar transporters. Each row of the heatmap is represented as condition – time point.

These evolutionary findings coupled with the higher expression of xylF on the microarray make this gene a strong candidate for further studies and industrial use.

## 4. Discussion

Gene duplication is an important phenomenon in eukaryotes as a source of novelty and adaptation [70], however in prokaryotes it is rarely seen, as arisen duplications are costly to maintain [1]. Thus, finding multiple copies of a gene for a given family may reflect adaptation in response to a new environmental pressure or functional specialization within the same environment [8]. A large proportion of gene family retractions were found in the *Thermoanaerobacterium* branch, a much higher retraction than observed in other species as shown in figure 1. This could be related to the stressful environments in which these organisms are found, as it is known that gene deletions can happen in stressful environments as a consequence of reduced usage, reducing energetic costs to the organism [3]. Even though this retraction was seen, Family OG55 of insertion sequences was found to be under rapid evolution.

Nucleotide substitutions that undergo positive selection through evolution on a given gene, as a consequence of an increase in the organism’s fitness, usually leave traces such as an altered rate of substitutions for an amino acid site. Attempting to correlate selection marks with industry desired phenotypes is a newer approach that may help to choose targets for bioengineering of industrial microorganisms [7,21,57,77]. Positive selection clues here revealed three gene families that seem to be important in xylose metabilosm. The OG55 transposase, found to be under positive selection, is positioned side-by-side with a xilose isomerase domain protein and an araC on the *T*.*sac* genome. Some studies have also shown that transposable elements may be beneficial by activating metabolism, resistance or acting as a defense mechanism [19,25].

Family OG963 was annotated as Succinate dehydrogenase, which is an oxidoreductase from the complex II electron transport chain. For the 3 genes in this family, an InterproScan search revealed an heterodisulfide reductase domain, which is associated with methanogenic reduction of ferredoxin [33], and in thermophilic bacteria, hydrogen is formed from ferredoxin, enabling extra ATP production [62]. One site in this region was found to be under positive selection, and AFK87395.1 was found to be more expressed in some xylose conditions, as seen in the microarray data. In *T. sac*, it was also previously described that Ferredoxin:NAD+ Oxidorreductase is essential for Ethanol formation [68].

Regarding families related to sugar metabolism, families OG1224, OG1421 and OG1052 showed a greater number of genes on our naïve approach. Family OG1224 of PH1107 glycosidase is related to degradation of glycoside bonds of carbohydrates, such as xylan and other complex sugars, which can help to explain its adaptations to ferment these carbohydrates directly, even more so with AFK87323.1 being upregulated on xylose conditions. Family OG1421 of acetoin dehydrogenase was particularly interesting as AFK86672.1 is upregulated in many xylose conditions as seen in the microarray data; as acetoin can be a fermentation inhibitor, this reaction could be extremely important for cofactor regeneration and disinhibition, especially in stressful conditions [78], furthermore, it has been reported that acetoin can also be used as a carbon source by *Bacillus subtilis* [29,76]. Regarding OG1052, in *T. sac*, it has been shown that PFOR is present as at least six clusters, with AFK87181.1 as a member of pforF and AFK86082.1 as a member of pforD. These genes are important for pyruvate dissimilation, with pforA (Tsac_0046/ AFK85084.1; OG320) as the most important member for ethanol production [85]. As pforA is not a member of OG1052, multiple copies of PFOR with slight differences may help alleviate redox needs for the cells and quicken reactions, as hinted by both the microarray data and STRINGdb network [65], which shows that both proteins are co-expressed with 4Fe-4S ferredoxin and other genes of this metabolical pathway.

As mentioned before, one of the bottlenecks in industrial 2G fermentation is inhibition of pentose transport by C6 sugars, thus, finding better xylose transporters is desirable to mitigate this occurrence.

Studies with MFS sugar transporters show that most are found to be under purifying selection [73,79], and better fermenting species are known to have more gene duplications, suggesting that sugar transport evolved in a multi-genic strategy to increase transport speed while maintaining their sequence relatively unchanged [44]. Because of the ATP dependency by ABC transporters, we hypothesized that adaptations should have occurred to compensate the risk of dispending energy to capture sugar molecules. Also, selective pressure signals should appear most likely on the sugar binding protein (xylF) as this energy expenditure would signify specificity to xylose. Thus, we searched for evidence of positive selection on families related to sugar transport, both MFS and ABC. No MFS transporters or xylFGH operon members had evidence of positive selection, except xylF (OG1742), which was found to be positively selected. This is interesting because xylF being the xylFGH operon’s extracellular protein, responsible for capturing xylose molecules, and being and ABC transporter, strongly indicates specificity to xylose to compensate the energy expenditure in transporting solutes. Moreover, most adaptations found in sugar transporters for yeast after rounds of directed evolution or mutagenesis are also in amino acid residues close to the predicted binding sites [20,56]. Evidence of positive selection, coupled with higher expression during xylose fermentation as seen on the published microarray suggests a higher adaptation for xylose metabolism in this organism by increasing its efficiency on capturing xylose and may be a key reason on why it is capable to co-ferment glucose and xylose. Not many sugar transporters specific for xylose are known and having an independent route for assimilation may alleviate competition of sugar molecules to the active site of promiscuous MFS transporters.

Regarding the possibility of using this transporter for present microbial cell factories, in eukaryotes there are almost no ABC transporters functioning on importing molecules, only exporters [75]. However, for industrial yeast, in addition to extracellular pumps related to drug removal and detoxification, there are some ABC importers, such as AUS1 and PDR11 which are related to external sterol assimilation for ergosterol production [35,38]. Moreover, horizontal gene transfer has been reported from bacteria to fungi [36,59], including some genes associated with xylose metabolism, such as Xylose Isomerase [50]. This indicates that adapting current *T. sac* xylFGH operon for use in yeast may be a possibility to help co-consumption of xylose and glucose during 2G ethanol fermentation, such as by mutating key residues in MFS transporters using *T. sac* xylF as a model.

*Thermoanaerobacter* species, such as *Thermoanaerobacterium saccharolyticum*, have several traits that can benefit industrial biotechnological fermentations, especially 2G ethanol production, for which *T. sac* has been engineered to metabolize at a great yield. In addition to genomic and transcriptomic resources already published, a deep evolutionary analysis and understanding of *T. sac* by comparing its genome against other bacteria groups is presented. Genomic adaptations to environmental stress are shown, such as heat stress and iron reduction, as well as xylose metabolism, seen at the specific xylFGH xylose operon, which has been also found in expression data, showing that evolutionary exploratory analysis can be useful for biotechnological prospecting. These data can serve as basis on searching for targets for industrial adaptation of *T. sac*, as well as reveal novel genes for yeast engineering.

## References

[1] Adler, M., Anjum, M., Berg, O.G., Andersson, D.I., Sandegren, L. (2014) High fitness costs and instability of gene duplications reduce rates of evolution of new genes by duplication-divergence mechanisms. Molecular Biology and Evolution 31(6), 1526–35, Doi: 10.1093/molbev/msu111.

[2] Ai, H., Zhang, J., Yang, M., Yu, P., Li, S., Zhu, M., Dong, H., Wang, S., Wang, J. (2014) Draft Genome Sequence of an Anaerobic, Thermophilic Bacterium, Thermoanaerobacterium aotearoense SCUT27, Isolated from a Hot Spring in China. Genome Announcements 2(1), Doi: 10.1128/genomea.00041-14.

[3] Albalat, R., Cañestro, C. (2016) Evolution by gene loss. Nat Rev Genet 17(7), 379–91, Doi: 10.1038/nrg.2016.39.

[4] Altermann, E., Russell, W.M., Azcarate-Peril, M.A., Barrangou, R., Buck, B.L., McAuliffe, O., Souther, N., Dobson, A., Duong, T., Callanan, M., Lick, S., Hamrick, A., Cano, R., Klaenhammer, T.R. (2005) Complete genome sequence of the probiotic lactic acid bacterium Lactobacillus acidophilus NCFM. Proceedings of the National Academy of Sciences of the United States of America 102(11), 3906–12, Doi: 10.1073/pnas.0409188102.

[5] Baesman, S.M., Stolz, J.F., Kulp, T.R., Oremland, R.S. (2009) Enrichment and isolation of Bacillus beveridgei sp. nov., a facultative anaerobic haloalkaliphile from Mono Lake, California, that respires oxyanions of tellurium, selenium, and arsenic. Extremophiles 13(4), 695–705, Doi: 10.1007/s00792-009-0257-z.

[6] Blattner, F.R., Plunkett, G., Bloch, C.A., Perna, N.T., Burland, V., Riley, M., Collado-Vides, J., Glasner, J.D., Rode, C.K., Mayhew, G.F., Gregor, J., Davis, N.W., Kirkpatrick, H.A., Goeden, M.A., Rose, D.J., Mau, B., Shao, Y. (1997) The complete genome sequence of Escherichia coli K-12. Science 277(5331), 1453–62, Doi: 10.1126/science.277.5331.1453.

[7] Borelli, G., Fiamenghi, M.B., Dos Santos, L.V., Carazzolle, M.F., Pereira, G.A.G., José, J., Zhang, G. (2019) Positive Selection Evidence in Xylose-Related Genes Suggests Methylglyoxal Reductase as a Target for the Improvement of Yeasts’ Fermentation in Industry. Genome Biology and Evolution 11(7), 1923–38, Doi: 10.1093/gbe/evz036.

[8] Bratlie, M.S., Johansen, J., Sherman, B.T., Huang, D.W., Lempicki, R.A., Drabløs, F. (2010) Gene duplications in prokaryotes can be associated with environmental adaptation. BMC Genomics 11(1), Doi: 10.1186/1471-2164-11-588.

[9] Brown, S.D., Begemann, M.B., Mormile, M.R., Wall, J.D., Han, C.S., Goodwin, L.A., Pitluck, S., Land, M.L., Hauser, L.J., Elias, D.A. (2011) Complete genome sequence of the haloalkaliphilic, hydrogen-producing bacterium Halanaerobium hydrogeniformans. Journal of Bacteriology 193(14), 3682–3, Doi: 10.1128/JB.05209-11.

[10] Buchfink, B., Xie, C., Huson, D.H. (2014) Fast and sensitive protein alignment using DIAMOND. Nature Methods 12(1), 59–60, Doi: 10.1038/nmeth.3176.

[11] Bueno, J.G.R., Borelli, G., Corrêa, T.L.R., Fiamenghi, M.B., José, J., de Carvalho, M., de Oliveira, L.C., Pereira, G.A.G., dos Santos, L.V. (2020) Novel xylose transporter Cs4130 expands the sugar uptake repertoire in recombinant Saccharomyces cerevisiae strains at high xylose concentrations. Biotechnology for Biofuels 13(1), 145, Doi: 10.1186/s13068-020-01782-0.

[12] Burroughs, A.M., Allen, K.N., Dunaway-Mariano, D., Aravind, L. (2006) Evolutionary Genomics of the HAD Superfamily: Understanding the Structural Adaptations and Catalytic Diversity in a Superfamily of Phosphoesterases and Allied Enzymes. Journal of Molecular Biology 361(5), 1003–34, Doi: 10.1016/j.jmb.2006.06.049.

[13] Cui, J., Davidson, A.L. (2011) ABC solute importers in bacteria. Essays in Biochemistry 50(1), 85–99, Doi: 10.1042/BSE0500085.

[14] Cui, J., Maloney, M.I., Olson, D.G., Lynd, L.R. (2020) Conversion of phosphoenolpyruvate to pyruvate in Thermoanaerobacterium saccharolyticum. Metabolic Engineering Communications 10, e00122, Doi: 10.1016/j.mec.2020.e00122.

[15] Currie, D.H., Raman, B., Gowen, C.M., Tschaplinski, T.J., Land, M.L., Brown, S.D., Covalla, S.F., Klingeman, D.M., Yang, Z.K., Engle, N.L., Johnson, C.M., Rodriguez, M., Joe Shaw, A., Kenealy, W.R., Lynd, L.R., Fong, S.S., Mielenz, J.R., Davison, B.H., Hogsett, D.A., Herring, C.D. (2015) Genome-scale resources for Thermoanaerobacterium saccharolyticum. BMC Systems Biology 9(1), 1–15, Doi: 10.1186/s12918-015-0159-x.

[16] De Bie, T., Cristianini, N., Demuth, J.P., Hahn, M.W. (2006) CAFE: A computational tool for the study of gene family evolution. Bioinformatics 22(10), 1269–71, Doi: 10.1093/bioinformatics/btl097.

[17] Dideberg, O., Charlier, P., Dive, G., Joris, B., Frère, J.M., Ghuysen, J.M. (1982) Structure of a Zn 2+ - containing D -alanyl-D -alanine-cleaving carboxypeptidase at 2.5 Å resolution. Nature 1982 299:5882 299(5882), 469–70, Doi: 10.1038/299469a0.

[18] Emms, D.M., Kelly, S. (2015) OrthoFinder: solving fundamental biases in whole genome comparisons dramatically improves orthogroup inference accuracy. Genome Biology 16(1), 157, Doi: 10.1186/s13059-015-0721-2.

[19] Fan, C., Wu, Y.H., Decker, C.M., Rohani, R., Gesell Salazar, M., Ye, H., Cui, Z., Schmidt, F., Huang, W.E. (2019) Defensive Function of Transposable Elements in Bacteria. ACS Synthetic Biology 8(9), 2141–51, Doi: 10.1021/acssynbio.9b00218.

[20] Farwick, A., Bruder, S., Schadeweg, V., Oreb, M., Boles, E. (2014) Engineering of yeast hexose transporters to transport D-xylose without inhibition by D-glucose. Proceedings of the National Academy of Sciences of the United States of America 111(14), 5159–64, Doi: 10.1073/pnas.1323464111.

[21] Fiamenghi, M.B., Bueno, J.G.R., Camargo, A.P., Borelli, G., Carazzolle, M.F., Pereira, G.A.G., dos Santos, L.V., José, J. (2022) Machine learning and comparative genomics approaches for the discovery of xylose transporters in yeast. Biotechnology for Biofuels and Bioproducts 15(1), 57, Doi: 10.1186/s13068-022-02153-7.

[22] Foster, S.J. (1991) Cloning, expression, sequence analysis and biochemical characterization of an autolytic amidase of Bacillus subtilis 168 trpC2. Microbiology 137(8), 1987–98, Doi: 10.1099/00221287-137-8-1987.

[23] Furdui, C., Ragsdale, S.W. (2000) The role of pyruvate ferredoxin oxidoreductase in pyruvate synthesis during autotrophic growth by the Wood-Ljungdahl pathway. Journal of Biological Chemistry 275(37), 28494–9, Doi: 10.1074/jbc.M003291200.

[24] Guo, X., Liao, Z., Holtzapple, M., Hu, Q., Zhao, B. (2017) Draft genome sequence of natranaerobius trueperi DSM 18760T, an anaerobic, halophilic, alkaliphilic, thermotolerant bacterium isolated from a soda lake. Genome Announcements 5(36), Doi: 10.1128/genomeA.00785-17.

[25] Hall, B.G. (1998) Activation of the bgl operon by adaptive mutation. Molecular Biology and Evolution 15(1), 1–5, Doi: 10.1093/oxfordjournals.molbev.a025842.

[26] Heider, J., Mai, X., Adams, M.W.W. (1996) Characterization of 2-ketoisovalerate ferredoxin oxidoreductase, a new and reversible coenzyme A-dependent enzyme involved in peptide fermentation by hyperthermophilic archaea. Journal of Bacteriology 178(3), 780–7, Doi: 10.1128/jb.178.3.780-787.1996.

[27] Hemme, C.L., Mouttaki, H., Lee, Y.J., Zhang, G., Goodwin, L., Lucas, S., Copeland, A., Lapidus, A., Del Rio, T.G., Tice, H., Saunders, E., Brettin, T., Detter, J.C., Han, C.S., Pitluck, S., Land, M.L., Hauser, L.J., Kyrpides, N., Mikhailova, N., He, Z., Wu, L., Van Nostrand, J.D., Henrissat, B., He, Q., Lawson, P.A., Tanner, R.S., Lynd, L.R., Wiegel, J., Fields, M.W., Arkin, A.P., Schadt, C.W., Stevenson, B.S., McInerney, M.J., Yang, Y., Dong, H., Xing, D., Ren, N., Wang, A., Huhnke, R.L., Mielenz, J.R., Ding, S.Y., Himmel, M.E., Taghavi, S., Van Der Lelie, D., Rubin, E.M., Zhou, J. (2010) Sequencing of multiple clostridial genomes related to biomass conversion and biofuel production. Journal of Bacteriology 192(24), 6494–6, Doi: 10.1128/JB.01064-10.

[28] Hsu, J.-L., Peng, H.-L., Chang, H.-Y. (2008) The ATP-binding motif in AcoK is required for regulation of acetoin catabolism in Klebsiella pneumoniae CG43. Biochemical and Biophysical Research Communications 376(1), 121–7, Doi: 10.1016/j.bbrc.2008.08.103.

[29] Huang, M., Oppermann-Sanio, F.B., Steinbüchel, A. (1999) Biochemical and Molecular Characterization of the Bacillus subtilis Acetoin Catabolic Pathway. Journal of Bacteriology 181(12), 3837–41, Doi: 10.1128/JB.181.12.3837-3841.1999.

[30] Huerta-Cepas, J., Forslund, K., Coelho, L.P., Szklarczyk, D., Jensen, L.J., von Mering, C., Bork, P. (2017) Fast Genome-Wide Functional Annotation through Orthology Assignment by eggNOG-Mapper. Molecular Biology and Evolution 34(8), 2115–22, Doi: 10.1093/molbev/msx148.

[31] Ivanova, N., Sikorski, J., Chertkov, O., Nolan, M., Lucas, S., Hammon, N., Deshpande, S., Cheng, J.F., Tapia, R., Han, C., Goodwin, L., Pitluck, S., Huntemann, M., Liolios, K., Pagani, I., Mavromatis, K., Ovchinikova, G., Pati, A., Chen, A., Palaniappan, K., Land, M., Hauser, L., Brambilla, E.M., Kannan, K.P., Rohde, M., Tindall, B.J., Göker, M., Detter, J.C., Woyke, T., Bristow, J., Eisen, J.A., Markowitz, V., Hugenholtz, P., Kyrpides, N.C., Klenk, H.P., Lapidus, A. (2011) Complete genome sequence of the extremely Halophilic Halanaerobium praevalens type strain (GSL T). Standards in Genomic Sciences 4(3), 312–21, Doi: 10.4056/sigs.1824509.

[32] Joe Shaw, A., Jenney, F.E., Adams, M.W.W.W., Lynd, L.R. (2008) End-product pathways in the xylose fermenting bacterium, Thermoanaerobacterium saccharolyticum. Enzyme and Microbial Technology 42(6), 453–8, Doi: 10.1016/j.enzmictec.2008.01.005.

[33] Kaster, A.K., Moll, J., Parey, K., Thauer, R.K. (2011) Coupling of ferredoxin and heterodisulfide reduction via electron bifurcation in hydrogenotrophic methanogenic archaea. Proceedings of the National Academy of Sciences of the United States of America 108(7), 2981–6, Doi: 10.1073/pnas.1016761108.

[34] Katoh, K., Standley, D.M. (2013) MAFFT Multiple Sequence Alignment Software Version 7: Improvements in Performance and Usability. Molecular Biology and Evolution 30(4), 772–80, Doi: 10.1093/molbev/mst010.

[35] Kohut, P., Wüstner, D., Hronska, L., Kuchler, K., Hapala, I., Valachovic, M. (2011) The role of ABC proteins Aus1p and Pdr11p in the uptake of external sterols in yeast: Dehydroergosterol fluorescence study. Biochemical and Biophysical Research Communications 404(1), 233–8, Doi: 10.1016/j.bbrc.2010.11.099.

[36] Kominek, J., Doering, D.T., Opulente, D.A., Shen, X.-X., Zhou, X., DeVirgilio, J., Hulfachor, A.B., Groenewald, M., Mcgee, M.A., Karlen, S.D., Kurtzman, C.P., Rokas, A., Hittinger, C.T. (2019) Eukaryotic Acquisition of a Bacterial Operon. Cell 176(6), 1356-1366.e10, Doi: 10.1016/j.cell.2019.01.034.

[37] Kück, P., Longo, G.C. (2014) FASconCAT-G: Extensive functions for multiple sequence alignment preparations concerning phylogenetic studies. Frontiers in Zoology 11(1), 81, Doi: 10.1186/s12983-014-0081-x.

[38] Kumari, S., Kumar, M., Gaur, N.A., Prasad, R. (2021) Multiple roles of ABC transporters in yeast. Fungal Genetics and Biology 150(January), 103550, Doi: 10.1016/j.fgb.2021.103550.

[39] Kunst, F., Ogasawara, N., Moszer, I., Albertini, A.M., Alloni, G., Azevedo, V., Bertero, M.G., Bessières, P., Bolotin, A., Borchert, S., Borriss, R., Boursier, L., Brans, A., Braun, M., Brignell, S.C., Bron, S., Brouillet, S., Bruschi, C.V., Caldwell, B., Capuano, V., Carter, N.M., Choi, S.K., Codani, J.J., Connerton, I.F., Cummings, N.J., Daniel, R.A., Denizot, F., Devine, K.M., Düsterhöft, A., Ehrlich, S.D., Emmerson, P.T., Entian, K.D., Errington, J., Fabret, C., Ferrari, E., Foulger, D., Fritz, C., Fujita, M., Fujita, Y., Fuma, S., Galizzi, A., Galleron, N., Ghim, S.Y., Glaser, P., Goffeau, A., Golightly, E.J., Grandi, G., Guiseppi, G., Guy, B.J., Haga, K., Haiech, J., Harwood, C.R., Hénaut, A., Hilbert, H., Holsappel, S., Hosono, S., Hullo, M.F., Itaya, M., Jones, L., Joris, B., Karamata, D., Kasahara, Y., Klaerr-Blanchard, M., Klein, C., Kobayashi, Y., Koetter, P., Koningstein, G., Krogh, S., Kumano, M., Kurita, K., Lapidus, A., Lardinois, S., Lauber, J., Lazarevic, V., Lee, S.M., Levine, A., Liu, H., Masuda, S., Mauël, C., Médigue, C., Medina, N., Mellado, R.P., Mizuno, M., Moestl, D., Nakai, S., Noback, M., Noone, D., O’Reilly, M., Ogawa, K., Ogiwara, A., Oudega, B., Park, S.H., Parro, V., Pohl, T.M., Portetelle, D., Porwollik, S., Prescott, A.M., Presecan, E., Pujic, P., Purnelle, B., Rapoport, G., Rey, M., Reynolds, S., Rieger, M., Rivolta, C., Rocha, E., Roche, B., Rose, M., Sadaie, Y., Sato, T., Scanlan, E., Schleich, S., Schroeter, R., Scoffone, F., Sekiguchi, J., Sekowska, A., Seror, S.J., Serror, P., Shin, B.S., Soldo, B., Sorokin, A., Tacconi, E., Takagi, T., Takahashi, H., Takemaru, K., Takeuchi, M., Tamakoshi, A., Tanaka, T., Terpstra, P., Tognoni, A., Tosato, V., Uchiyama, S., Vandenbol, M., Vannier, F., Vassarotti, A., Viari, A., Wambutt, R., Wedler, E., Wedler, H., Weitzenegger, T., Winters, P., Wipat, A., Yamamoto, H., Yamane, K., Yasumoto, K., Yata, K., Yoshida, K., Yoshikawa, H.F., Zumstein, E., Yoshikawa, H., Danchin, A. (1997) The complete genome sequence of the gram-positive bacterium Bacillus subtilis. Nature 390(6657), 249–56, Doi: 10.1038/36786.

[40] Kwak, S., Jin, Y.-S. (2017) Production of fuels and chemicals from xylose by engineered Saccharomyces cerevisiae: a review and perspective. Microbial Cell Factories 16(1), 82, Doi: 10.1186/s12934-017-0694-9.

[41] Lee, Y.E., Jain, M.K., Lee, C., Lowe, S.E., Zeikus, J.G. (1993) Taxonomic distinction of saccharolytic thermophilic anaerobes. International Journal of Systematic Bacteriology 43(1), 41–51, Doi: 10.1099/00207713-43-1-41.

[42] Lewinson, O., Livnat-Levanon, N. (2017) Mechanism of Action of ABC Importers: Conservation, Divergence, and Physiological Adaptations. Journal of Molecular Biology 429(5), 606–19, Doi: 10.1016/j.jmb.2017.01.010.

[43] Lin, L., Song, H., Tu, Q., Qin, Y., Zhou, A., Liu, W., He, Z., Zhou, J., Xu, J. (2011) The Thermoanaerobacter glycobiome reveals mechanisms of pentose and hexose co-utilization in bacteria. PLoS Genetics 7(10), e1002318, Doi: 10.1371/journal.pgen.1002318.

[44] Lin, Z., Li, W.H. (2011) Expansion of Hexose Transporter Genes Was Associated with the Evolution of Aerobic Fermentation in Yeasts. Molecular Biology and Evolution 28(1), 131–42, Doi: 10.1093/MOLBEV/MSQ184.

[45] Liu, S.-Y., Rainey, F.A., Morgan, H.W., Mayer, F., Wiegel, J. (1996) Thermoanaerobacterium aotearoense sp. nov., a Slightly Acidophilic, Anaerobic Thermophile Isolated from Various Hot Springs in New Zealand, and Emendation of the Genus Thermoanaerobacterium. International Journal of Systematic Bacteriology 46(2), 388–96, Doi: 10.1099/00207713-46-2-388.

[46] Mai, V., Lorenz, W.W., Wiegel, J. (2006) Transformation of Thermoanaerobacterium sp. strain JW/SL-YS485 with plasmid pIKM1 conferring kanamycin resistance. FEMS Microbiology Letters 148(2), 163–7, Doi: 10.1111/j.1574-6968.1997.tb10283.x.

[47] Mai, V., Wiegel, J. (2000) Advances in development of a genetic system for Thermoanaerobacterium spp.: Expression of genes encoding hydrolytic enzymes, development of a second shuttle vector, and integration of genes into the chromosome. Applied and Environmental Microbiology 66(11), 4817–21, Doi: 10.1128/AEM.66.11.4817-4821.2000.

[48] Mead, D., Drinkwater, C., Brumm, P.J. (2013) Genomic and Enzymatic Results Show Bacillus cellulosilyticus Uses a Novel Set of LPXTA Carbohydrases to Hydrolyze Polysaccharides. PLoS ONE 8(4), e61131, Doi: 10.1371/journal.pone.0061131.

[49] Morita, H., Hidehiro, T.O.H., Fukuda, S., Horikawa, H., Oshima, K., Suzuki, T., Murakami, M., Hisamatsu, S., Kato, Y., Takizawa, T., Fukuoka, H., Yoshimura, T., Itoh, K., O’Sullivan, D.J., Mckay, L.L., Ohno, H., Kikuchi, J., Masaoka, T., Hattori, M. (2008) Comparative genome analysis of Lactobacillus renteri and Lactobacillus fermentum reveal a genomic Island for reuterin and cobalamin production. DNA Research 15(3), 151–61, Doi: 10.1093/dnares/dsn009.

[50] Murphy, C.L., Youssef, N.H., Hanafy, R.A., Couger, M.B., Stajich, J.E., Wang, Y., Baker, K., Dagar, S.S., Griffith, G.W., Farag, I.F., Callaghan, T.M., Elshahed, M.S. (2019) Horizontal Gene Transfer as an Indispensable Driver for Evolution of Neocallimastigomycota into a Distinct Gut-Dwelling Fungal Lineage. Applied and Environmental Microbiology 85(15), e00988–19, Doi: 10.1128/AEM.00988-19.

[51] Murrell, B., Wertheim, J.O., Moola, S., Weighill, T., Scheffler, K. (2012) Detecting Individual Sites Subject to Episodic Diversifying Selection. PLoS Genet 8(7), 1002764, Doi: 10.1371/journal.pgen.1002764.

[52] Nölling, J., Breton, G., Omelchenko, M.V., Makarova, K.S., Zeng, Q., Gibson, G., Hong Mei Lee, Dubois, J., Qiu, D., Hitti, J., Aldredge, T., Ayers, M., Bashirzadeh, R., Bochner, H., Boivin, M., Bross, S., Bush, D., Butler, C., Caron, A., Caruso, A., Cook, R., Daggett, P., Deloughery, C., Egan, J., Ellston, D., Engelstein, M., Ezedi, J., Gilbert, K., Goyal, A., Guerin, J., Ho, T., Holtham, K., Joseph, P., Keagle, P., Kozlovsky, J., LaPlante, M., LeBlanc, G., Lumm, W., Majeski, A., McDougall, S., Mank, P., Mao, J.I., Nocco, D., Patwell, D., Phillips, J., Pothier, B., Prabhakar, S., Richterich, P., Rice, P., Rosetti, D., Rossetti, M., Rubenfield, M., Sachdeva, M., Snell, P., Spadafora, R., Spitzer, L., Shimer, G., Thomann, H.U., Vicaire, R., Wall, K., Wang, Y., Weinstock, K., Lai Peng Wong, Wonsey, A., Xu, Q., Zhang, L., Wolf, Y.I., Tatusov, R.L., Sabathe, F., Doucette-Stamm, L., Soucaille, P., Daly, M.J., Bennett, G.N., Koonin, E.V., Smith, D.R. (2001) Genome sequence and comparative analysis of the solvent-producing bacterium Clostridium acetobutylicum. Journal of Bacteriology 183(16), 4823–38, Doi: 10.1128/JB.183.16.4823-4838.2001.

[53] Parkhill, J., Dougan, G., James, K.D., Thomson, N.R., Pickard, D., Wain, J., Churcher, C., Mungall, K.L., Bentley, S.D., Holden, M.T.G., Sebaihia, M., Baker, S., Basham, D., Brooks, K., Chillingworth, T., Connerton, P., Cronin, A., Davis, P., Davies, R.M., Dowd, L., White, N., Farrar, J., Feltwell, T., Hamlin, N., Haque, A., Hien, T.T., Holroyd, S., Jagels, K., Krogh, A., Larsen, T.S., Leather, S., Moule, S., Ó’Gaora, P., Parry, C., Quail, M., Rutherford, K., Simmonds, M., Skelton, J., Stevens, K., Whitehead, S., Barrell, B.G. (2001) Complete genome sequence of a multiple drug resistant Salmonella enterica serovar Typhi CT18. Nature 413(6858), 848–52, Doi: 10.1038/35101607.

[54] Parks, D.H., Chuvochina, M., Chaumeil, P.A., Rinke, C., Mussig, A.J., Hugenholtz, P. (2020) A complete domain-to-species taxonomy for Bacteria and Archaea. Nature Biotechnology 38(9), 1079–86, Doi: 10.1038/s41587-020-0501-8.

[55] Peng, H.L., Yang, Y.H., Deng, W.L., Chang, H.Y. (1997) Identification and characterization of acoK, a regulatory gene of the Klebsiella pneumoniae acoABCD operon. Journal of Bacteriology 179(5), 1497–504, Doi: 10.1128/jb.179.5.1497-1504.1997.

[56] Reider Apel, A., Ouellet, M., Szmidt-Middleton, H., Keasling, J.D., Mukhopadhyay, A. (2016) Evolved hexose transporter enhances xylose uptake and glucose/xylose co-utilization in Saccharomyces cerevisiae. Scientific Reports 6(1), 19512, Doi: 10.1038/srep19512.

[57] Riley, R., Haridas, S., Wolfe, K.H., Lopes, M.R., Hittinger, C.T., Göker, M., Salamov, A.A., Wisecaver, J.H., Long, T.M., Calvey, C.H., Aerts, A.L., Barry, K.W., Choi, C., Clum, A., Coughlan, A.Y., Deshpande, S., Douglass, A.P., Hanson, S.J., Klenk, H.-P., LaButti, K.M., Lapidus, A., Lindquist, E.A., Lipzen, A.M., Meier-Kolthoff, J.P., Ohm, R.A., Otillar, R.P., Pangilinan, J.L., Peng, Y., Rokas, A., Rosa, C.A., Scheuner, C., Sibirny, A.A., Slot, J.C., Stielow, J.B., Sun, H., Kurtzman, C.P., Blackwell, M., Grigoriev, I.V., Jeffries, T.W. (2016) Comparative genomics of biotechnologically important yeasts. Proceedings of the National Academy of Sciences 113(35), 9882–7, Doi: 10.1073/pnas.1603941113.

[58] Ronquist, F., Teslenko, M., van der Mark, P., Ayres, D.L., Darling, A., Höhna, S., Larget, B., Liu, L., Suchard, M.A., Huelsenbeck, J.P. (2012) MrBayes 3.2: efficient Bayesian phylogenetic inference and model choice across a large model space. Systematic Biology 61(3), 539–42, Doi: 10.1093/sysbio/sys029.

[59] Rosewich, U.L., Kistler, H.C. (2000) Role of Horizontal Gene Transfer in the Evolution of Fungi. Annu. Rev. Phytopathol. 38(1), 325–63, Doi: 10.1146/annurev.phyto.38.1.325.

[60] Rychener, L., Albon, S.I., Djordjevic, S.P., Chowdhury, P.R., Ziech, R.E., de Vargas, A.C., Frey, J., Falquet, L. (2017) Clostridium chauvoei, an evolutionary dead-end pathogen. Frontiers in Microbiology 8(JUN), 1054, Doi: 10.3389/fmicb.2017.01054.

[61] Sb, H., T, D., Mk, W., F, C. (2020) Modulation of Peptidoglycan Synthesis by Recycled Cell Wall Tetrapeptides. Cell Reports 31(4), Doi: 10.1016/J.CELREP.2020.107578.

[62] Shaw, A.J., Hogsett, D.A., Lynd, L.R. (2009) Identification of the [FeFe]-hydrogenase responsible for hydrogen generation in Thermoanaerobacterium saccharolyticum and demonstration of increased ethanol yield via hydrogenase knockout. Journal of Bacteriology 191(20), 6457–64, Doi: 10.1128/JB.00497-09.

[63] Shaw, A.J., Podkaminer, K.K., Desai, S.G., Bardsley, J.S., Rogers, S.R., Thorne, P.G., Hogsett, D.A., Lynd, L.R. (2008) Metabolic engineering of a thermophilic bacterium to produce ethanol at high yield. Proceedings of the National Academy of Sciences 105(37), 13769–74, Doi: 10.1073/pnas.0801266105.

[64] Smith, T.J., Hill, K.K., Foley, B.T., Detter, J.C., Munk, A.C., Bruce, D.C., Doggett, N.A., Smith, L.A., Marks, J.D., Xie, G., Brettin, T.S. (2007) Analysis of the neurotoxin complex genes in Clostridium botulinum A1-A4 and B1 strains: BoNT/A3, /Ba4 and /B1 clusters are located within plasmids. PLoS ONE 2(12), Doi: 10.1371/journal.pone.0001271.

[65] Szklarczyk, D., Gable, A.L., Lyon, D., Junge, A., Wyder, S., Huerta-Cepas, J., Simonovic, M., Doncheva, N.T., Morris, J.H., Bork, P., Jensen, L.J., Von Mering, C. (2019) STRING v11: Protein-protein association networks with increased coverage, supporting functional discovery in genome-wide experimental datasets. Nucleic Acids Research 47(D1), D607–13, Doi: 10.1093/nar/gky1131.

[66] Tan, M.-F., Gao, T., Liu, W.-Q., Zhang, C.-Y., Yang, X., Zhu, J.-W., Teng, M.-Y., Li, L., Zhou, R. (2015) MsmK, an ATPase, Contributes to Utilization of Multiple Carbohydrates and Host Colonization of Streptococcus suis. PLOS ONE 10(7), e0130792, Doi: 10.1371/journal.pone.0130792.

[67] Tanaka, K.J., Song, S., Mason, K., Pinkett, H.W. (2018) Selective substrate uptake: The role of ATP-binding cassette (ABC) importers in pathogenesis. Biochimica et Biophysica Acta (BBA) -Biomembranes 1860(4), 868–77, Doi: 10.1016/j.bbamem.2017.08.011.

[68] Tian, L., Lo, J., Shao, X., Zheng, T., Olson, D.G., Lynd, L.R. (2016) Ferredoxin: NAD+ oxidoreductase of Thermoanaerobacterium saccharolyticum and its role in ethanol formation. Applied and Environmental Microbiology 82(24), 7134–41, Doi: 10.1128/AEM.02130-16.

[69] Törönen, P., Medlar, A., Holm, L. (2018) PANNZER2: A rapid functional annotation web server. Nucleic Acids Research 46(W1), W84–8, Doi: 10.1093/nar/gky350.

[70] Treangen, T.J., Rocha, E.P.C. (2011) Horizontal transfer, not duplication, drives the expansion of protein families in prokaryotes. PLoS Genetics 7(1), e1001284, Doi: 10.1371/journal.pgen.1001284.

[71] Trifinopoulos, J., Nguyen, L.-T., von Haeseler, A., Minh, B.Q. (2016) W-IQ-TREE: a fast online phylogenetic tool for maximum likelihood analysis. Nucleic Acids Research 44(18), 1–4, Doi: 10.1093/nar/gkw256.

[72] Tsakraklides, V., Shaw, A.J., Miller, B.B., Hogsett, D.A., Herring, C.D. (2012) Carbon catabolite repression in Thermoanaerobacterium saccharolyticum. Biotechnology for Biofuels 5(1), 85, Doi: 10.1186/1754-6834-5-85.

[73] Wang, W., Zhou, H., Ma, B., Owiti, A., Korban, S.S., Han, Y. (2016) Divergent Evolutionary Pattern of Sugar Transporter Genes is Associated with the Difference in Sugar Accumulation between Grasses and Eudicots. Sci Rep 6(1), 29153, Doi: 10.1038/srep29153.

[74] Webb, A.J., Homer, K.A., Hosie, A.H.F. (2008) Two closely related ABC transporters in Streptococcus mutans are involved in disaccharide and/or oligosaccharide uptake. Journal of Bacteriology 190(1), 168–78, Doi: 10.1128/JB.01509-07.

[75] Wilkens, S. (2015) Structure and mechanism of ABC transporters. F1000Prime Reports 7, Doi: 10.12703/P7-14.

[76] Williams, O.B., Morrow, M.B. (1928) The bacterial destruction of acetyl-methyl-carbinol. Journal of Bacteriology 16(1), 43–8, Doi: 10.1128/jb.16.1.43-48.1928.

[77] Wohlbach, D.J., Kuo, A., Sato, T.K., Potts, K.M., Salamov, A.A., Labutti, K.M., Sun, H., Clum, A., Pangilinan, J.L., Lindquist, E.A., Lucas, S., Lapidus, A., Jin, M., Gunawan, C., Balan, V., Dale, B.E., Jeffries, T.W., Zinkel, R., Barry, K.W., Grigoriev, I.V., Gasch, A.P. (2011) Comparative genomics of xylose-fermenting fungi for enhanced biofuel production. Proceedings of the National Academy of Sciences of the United States of America 108(32), 13212–7, Doi: 10.1073/pnas.1103039108.

[78] Xiao, Z., Xu, P. (2007) Acetoin metabolism in bacteria. Critical Reviews in Microbiology 33(2), 127–40, Doi: 10.1080/10408410701364604.

[79] Xu, X., Zeng, W., Li, Z., Wang, Z., Luo, Z., Li, J., Li, X., Yang, J. (2022) Genome-wide identification and expression profiling of sugar transporter genes in tobacco. Gene 835, 146652, Doi: 10.1016/j.gene.2022.146652.

[80] Yang, Z. (2007) PAML 4: Phylogenetic Analysis by Maximum Likelihood. Molecular Biology and Evolution 24(8), 1586–91, Doi: 10.1093/molbev/msm088.

[81] Zhang, J., Nielsen, R., Yang, Z. (2005) Evaluation of an Improved Branch-Site Likelihood Method for Detecting Positive Selection at the Molecular Level. Molecular Biology and Evolution 22(12), 2472–9, Doi: 10.1093/molbev/msi237.

[82] Zhao, B., Mesbah, N.M., Dalin, E., Goodwin, L., Nolan, M., Pitluck, S., Chertkov, O., Brettin, T.S., Han, J., Larimer, F.W., Land, M.L., Hauser, L., Kyrpides, N., Wiegel, J. (2011) Complete genome sequence of the anaerobic, halophilic alkalithermophile Natranaerobius thermophilus JW/NM-WN-LF. Journal of Bacteriology 193(15), 4023–4, Doi: 10.1128/JB.05157-11.

[83] Zhao, Z., Xian, M., Liu, M., Zhao, G. (2020) Biochemical routes for uptake and conversion of xylose by microorganisms. Biotechnology for Biofuels 13(1), 21, Doi: 10.1186/s13068-020-1662-x.

[84] Zheng, T., Lanahan, A.A., Lynd, L.R., Olson, D.G. (2018) The redox-sensing protein Rex modulates ethanol production in Thermoanaerobacterium saccharolyticum. PLOS ONE 13(4), e0195143, Doi: 10.1371/journal.pone.0195143.

[85] Zhou, J., Olson, D.G., Lanahan, A.A., Tian, L., Murphy, S.J.L., Lo, J., Lynd, L.R. (2015) Physiological roles of pyruvate ferredoxin oxidoreductase and pyruvate formate-lyase in Thermoanaerobacterium saccharolyticum JW/SL-YS485. Biotechnology for Biofuels 8(1), 138, Doi: 10.1186/s13068-015-0304-1.

