## Supplementary Figures for "Comparative Genomics of Firmicutes reveals probable adaptations for xylose fermentation in Thermoanaerobacterium saccharolyticum"

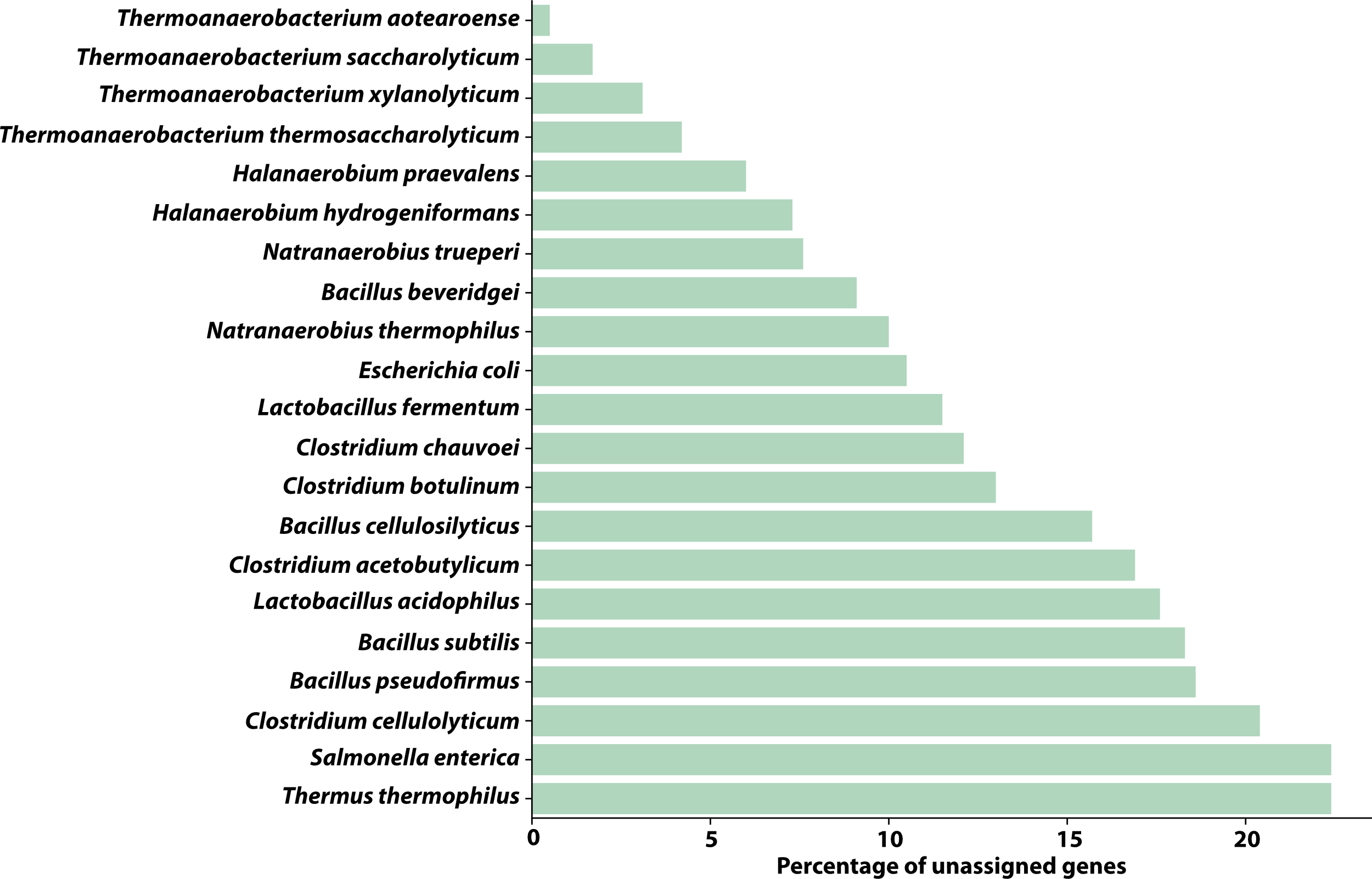

Supplementary figure 1. Percentage of unassigned genes per species as found by Orthofinder analysis

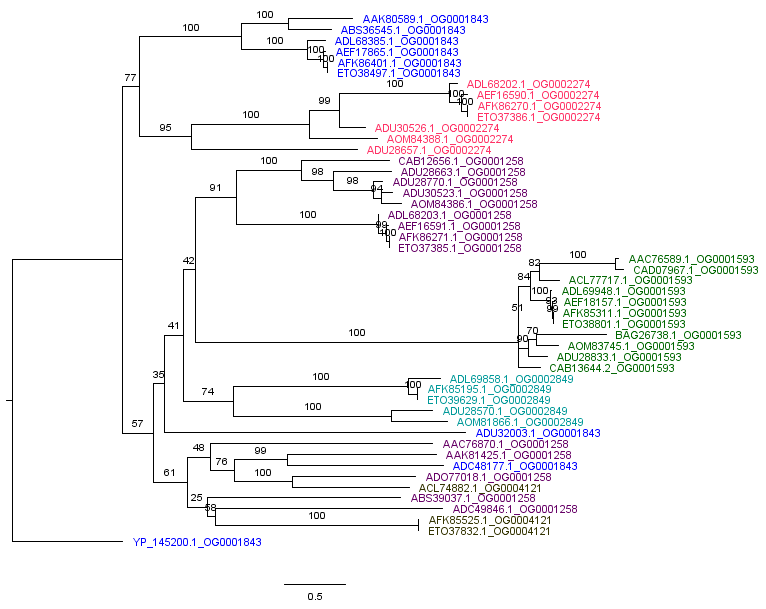

Supplementary fig2. Concatenated alignment of all Xylose Isomerase families
